# Neural Dynamics During the Generation and Evaluation of Creative and Non-Creative Ideas

**DOI:** 10.1101/2024.04.15.589621

**Authors:** Yoed N. Kenett, Evangelia G. Chrysikou, Dani S. Bassett, Sharon L. Thompson-Schill

## Abstract

What are the neural dynamics that drive creative thinking? Recent studies have provided much insight into the neural mechanisms of creative thought. Specifically, the interaction between the executive control, default mode, and salience brain networks has been shown to be an important marker of individual differences in creative ability. However, how these different brain systems might be recruited dynamically during the two key components of the creative process— generation and evaluation of ideas—remains far from understood. In the current study we applied state-of-the-art network neuroscience methodologies to examine the neural dynamics related to the generation and evaluation of creative and non-creative ideas using a novel within-subjects design. Participants completed two functional magnetic resonance imaging sessions, taking place a week apart. In the first imaging session, participants generated either creative (alternative uses) or non-creative (common characteristics) responses to common objects. In the second imaging session, participants evaluated their own creative and non-creative responses to the same objects. Network neuroscience methods were applied to examine and directly compare reconfiguration, integration, and recruitment of brain networks during these four conditions. We found that generating creative ideas led to significantly higher network reconfiguration than generating non-creative ideas, whereas evaluating creative and non-creative ideas led to similar levels of network integration. Furthermore, we found that these differences were attributable to different dynamic patterns of neural activity across the executive control, default mode, and salience networks. This study is the first to show within-subject differences in neural dynamics related to generating and evaluating creative and non-creative ideas.

## Introduction

Creativity involves multiple cognitive processes that allow for the generation of ideas that are both novel and useful ^1–4^. Recent research on the neural mechanisms of creative thinking has focused on how large-scale brain network connections may account for individual differences in creativity ^5,6^. The generation of creative ideas has been proposed to result from complex interactions between the executive and default mode networks, engagement of which appears to be mediated by activity within regions of the salience network during creative thinking ^7–12^. Indeed, increased resting-state functional brain connectivity between the inferior frontal cortex and key areas within the default mode network (DMN) has been associated with higher creative performance ^7,13^. Moreover, temporal connectivity between the default and salience (SN) networks has been shown to characterize performance earlier during a creative generation task, whereas connectivity between the default and executive (EN) networks characterizes performance later in the task ^7^.

Consistent with these findings, increased activity of DMN regions and the ventral anterior cingulate cortex (vACC) has been linked to the generation of original ideas; further, increased connectivity between the vACC and occipital-temporal areas was observed in participants who generated more original ideas ^14^. Studies have also consistently implicated the role of the hippocampus in creativity, which builds on prior knowledge that can then be recombined and utilized to create new and original ideas ^15–19^. Further, a recent study applied dynamic causal modeling to fMRI data and showed that prefrontal regions within the EN unidirectionally control posterior temporal and parietal regions of the DMN during divergent thinking ^20^. These results suggest that dynamic fluctuations of neural activity during creativity tasks within regions implicated in focused internal attention, cognitive control, and spontaneous thought may account for much individual variation in creative ability ^21–28^.

Despite the examination of such brain network interactions, past studies have tended to operationalize creativity through tasks that prioritize idea generation. However, the creative process has long been discussed as comprising alternating generative and evaluative components ^29,30^, with short– and long-term iterations between these phases being thought to occur many times during the performance of creativity tasks ^8,31^. Yet, the dynamics of these two phases—for example, whether they are serial or parallel—and the neural mechanisms that support such dynamics remain unclear.

Only a handful of studies have explored the neural bases of this twofold process of generating creative ideas while assessing their usefulness during creative ideation. Notably, Ellamil and colleagues ^32^ alternated participants’ generative and evaluative processes during a drawing task under fMRI to show that creative generation preferentially engaged DMN regions, whereas creative idea evaluation engaged both DMN and EN regions, as well as regions within the salience network. In line with these findings, reductions in the activity of left temporoparietal regions during participants’ evaluation of others’ creative ideas predicted higher creativity ratings, highlighting the importance of this region in evaluating—but also possibly inhibiting—creativity^33^.

Although this prior work points to interactions between the DMN and EN under a twofold model of creative cognition, the dynamic recruitment of different brain regions within these systems during the generation and evaluation of creative ideas, as well as any salience network mediation in these processes, remains poorly understood. A recent study leveraging temporal variability in cortical and cerebellar resting-state functional connectivity revealed that the dynamic reconfiguration of DMN and EN networks was associated with higher verbal creativity in a large sample of participants ^27^. Additionally, it has recently been shown that higher-creative participants show increased global and regional neural reconfiguration within EN and DMN regions during a creative relative to a non-creative task ^34^. Nevertheless, no prior study has examined the dynamic reconfiguration of large-scale brain networks during the generation and evaluation of *one’s own* creative ideas.

Here, we used network neuroscience approaches to examine the neural dynamics related to generation and evaluation of creative and non-creative ideas in a novel within-subjects design. We focused on three network neuroscience measures that capture such neural dynamics: Neural reconfiguration, recruitment, and integration ^35–40^. Neural reconfiguration quantifies how brain regions dynamically reconfigure their functional community across time and has been linked to neural dynamics in cognitive tasks such as learning, working memory, and linguistic processing ^34–36,41^. Neural integration reflects how brain regions from a specific neural system are functionally integrated with brain regions from other neural systems. Neural recruitment captures how brain regions connected with each other to form a neural system are further connected with other neural systems. These measures have been linked to variability in performance in various cognitive tasks, and allow investigators to examine how neural systems are integrated and/or recruited for specific cognitive tasks ^37^.

Although these measures have not been extensively used in the creativity literature (except for Ref. ^34^), they hold potential for understanding the complex processes implicated in creative thinking because they are not simply averaging functional connectivity across the brain. Rather, they are more sensitive to synchronous activity fluctuations across networks of regions, which may provide a better glimpse of the timing parameters and neural fluctuations of regional involvement during the creative process. Importantly, they uniquely allow us to test recent network neuroscience theories about the complexity of creative thinking, by going beyond correlating static functional connectivity patterns of activation with behavioral measures ^6,7,21^. Together, these three measures provide a quantitative approach toward examining the DMN, EN, and SN’s complex dynamical contributions to creative thinking.

Participants completed two functional magnetic resonance imaging sessions, taking place a week apart. In the first imaging session, participants generated either creative (alternative uses; AU) or non-creative (common characteristics; CC) responses to pictures of common objects. In the second imaging session, they evaluated their own creative and non-creative responses to the same objects. In addition—to account for possible differential processing in evaluating one’s own versus others’ ideas—participants evaluated a sample of creative and non-creative responses generated to different objects by an independent sample of participants.

Following pre-processing of fMRI data and network construction ^40,42^, we composed a dynamical functional brain network that represents neural reconfiguration, recruitment, and integration during the generation and evaluation of creative and non-creative ideas. We used dynamic community detection techniques ^43^ to extract groups of brain regions that were functionally connected to one another. We then characterized how these networks reconfigured, and how they were recruited and integrated over task conditions ^35–37^. Finally, to evaluate the validity of our findings, we compared these network dynamics to similar measures computed from participants’ resting-state fMRI data.

If creativity relies on the dynamic interactions among DMN and EN regions presumed to underly self-generated and goal-directed thought, we predicted that neural reconfiguration would be more pronounced during the generation—but not the evaluation—of creative ideas relative to the generation of common characteristics for objects. Prior work has suggested that areas with high reconfiguration become more significant for a behavior, because they participate in more neural processes across the brain ^36^. Higher reconfiguration would reflect the tendency of DMN and EN regions to temporarily change their community assignments and become transiently unstable to support maximum flexibility in creative idea generation ^36^. If creative thinking involves dynamic changes in the connectivity patterns among the DMN, EN, and SN during the creative process, we predicted higher integration of different neural systems across the creative process ^7^: higher integration of DMN regions with the SN in the generation stage, and higher DMN and EN integration in the evaluation stage of creative idea generation. Finally, if creative thinking involves recruiting cognitive systems at different stages of the creative process, such as cognitive control for idea evaluation and inhibition ^9,44^, we predicted higher recruitment specific to the EN during evaluation, and not generation, of creative ideas. Based on past research ^37,40^, we anticipated that recruitment and integration measures would be more pronounced during the evaluation relative to the generation of creative ideas due to the prioritization of comparisons between one’s responses and the task goals in the context of one’s past experience.

## Results

Our analysis process was as follows (**Fig. 1**). First, we recorded BOLD signals while participants generated and evaluated creative alternative uses (AU) and non-creative common characteristics (CC) ideas to common objects (**Fig. 1A**). Then using wavelet coherence analysis, we computed functional connectivity adjacency matrices for each AU and CC trial (**Fig. 1B**). We then pooled all condition-specific trials (Generation/Evaluation × AU/CC) and coupled them as a condition-specific multilayer network. Next, we applied a multilayer community detection approach to assign each brain region in each layer in each multilayer network to a community (**Fig. 1C**). Finally, we computed for each brain region its reconfiguration (the extent to which it changes its community assignment across layers; **Fig. 1D**), integration (the extent to which it integrates with brain regions from other neural systems; **Fig. 1E),** and recruitment (the extent to which it is recruited along with the whole neural system it belongs to in synchrony with other whole neural systems; **Fig. 1F**). We then averaged these three measures at the whole– and system-brain levels.

**Fig. 1.**
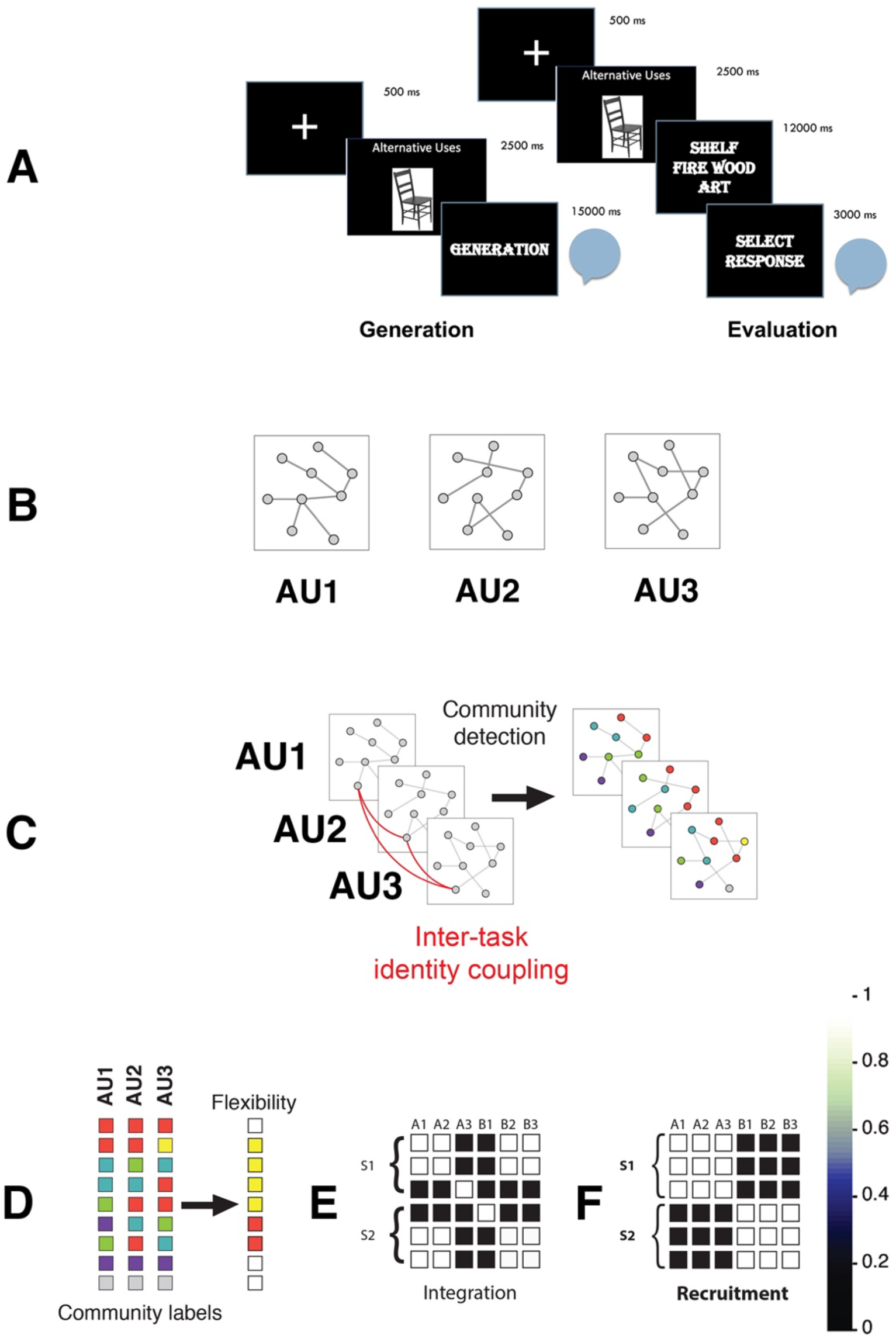
Illustration of our fMRI analysis pipeline. **(A)** BOLD signals were recorded while participants generated and evaluated creative (AU) and non-creative (CC) uses for common objects. **(B**) Using wavelet coherence analysis, functional connectivity adjacency matrices were computed for each AU and CC trial. **(C)** Trial level adjacency matrices were coupled together as a condition-specific multilayer (Generation/Evaluation × AU/CC) network. A multilayer network community detection approach was applied to assign each brain region in each layer in each multilayer network to a community. **(D)** The neural reconfiguration score of each brain region was computed. Each column represents a trial specific layer in the multilayer network, each square represents different brain regions, and colors represent different community assignments. Reconfiguration was computed as change in community assignment across layers. **(E)** Neural integration measures how brain regions from a specific neural system are functionally integrated with brain regions from other neural systems. **(F)** Neural recruitment captures how brain regions connected with each other to form a neural system are further connected with other neural systems. The illustration presents six brain regions (A1-3 and B1-3) related to two different neural systems (S1 and S2). The color bar represents the level of functional interaction between these brain regions across the two systems, according to neural integration and recruitment measures.

### Behavioral performance analysis

We first analyzed participants’ performance in the Generation-Evaluation task. In line with standard approaches used in creativity research ^45^, we measured participants fluency and obtained creativity scores of their responses to the AU and CC conditions. *Fluency* was measured as the number of responses that participants generated in 15 seconds to the objects presented to them in the Generation task (see *Methods*). *Creativity* of participants’ responses was measured as the quantitative semantic distance between an open-ended response and its prompt object, computed via computational modelling on textual corpora (SemDis; ^46^). The higher this score for an open-ended response is, the more original it is (see *Methods*). For each participant, their fluency and creativity scores were averaged across all objects separately for the AU and CC conditions. A paired-samples *t*-test on participants’ fluency scores revealed significantly higher fluency in generating CC responses (*M* = 3.61, *SD* = .79) than generating AU responses (*M* = 2.31, *SD* = .64), *t*(41) = 8.96, *p* < .001, *d* = .94. A paired-samples *t*-test on participants’ creativity scores revealed significantly lower creativity in the CC responses (*M* = .92, *SD* = .03) relative to the AU responses (*M* = .94, *SD* = .03), *t*(41) = –2.81, *p* = .008, *d* = .43. Thus, although participants were generating less AU responses during the AU than the CC task, their AU responses, as expected, were more creative (**Fig. 2**).

**Fig. 2.**
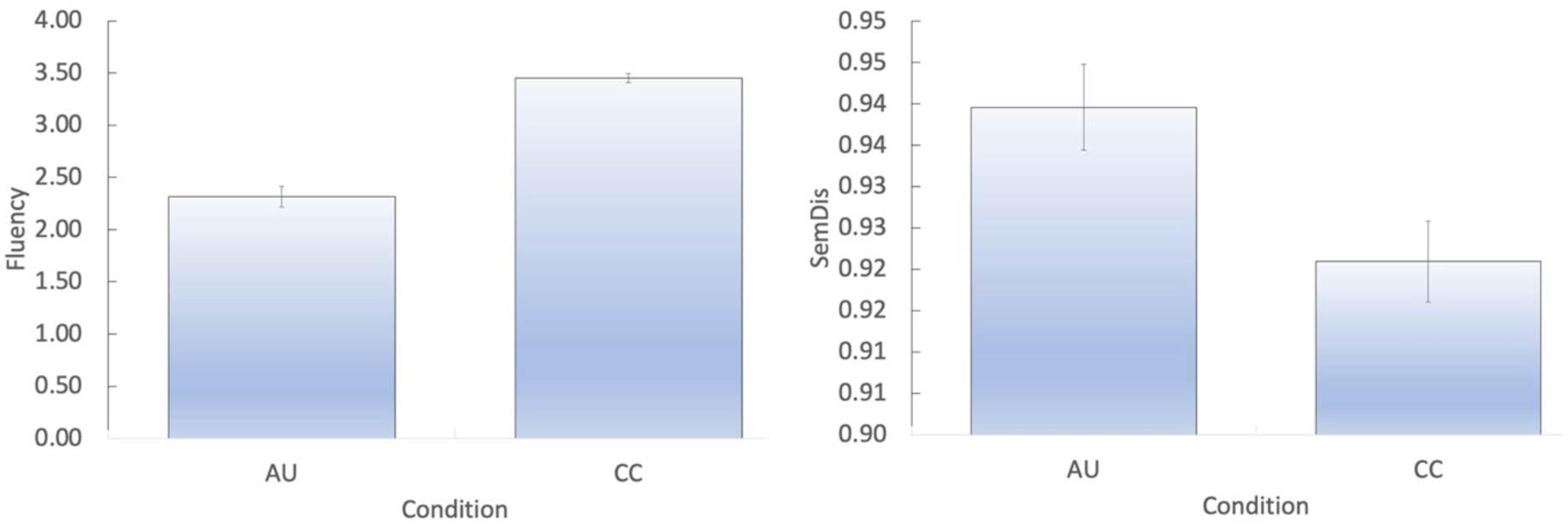
Behavioral analysis of participants’ responses in the AU and CC task: Left panel—fluency (number of responses); right panel—creativity (SemDis scores).

### Whole brain neural analysis

Next, we examined any possible differences in the whole-brain neural reconfiguration, integration, and recruitment measures across the four conditions (**Fig. 3)**.

**Fig. 3.**
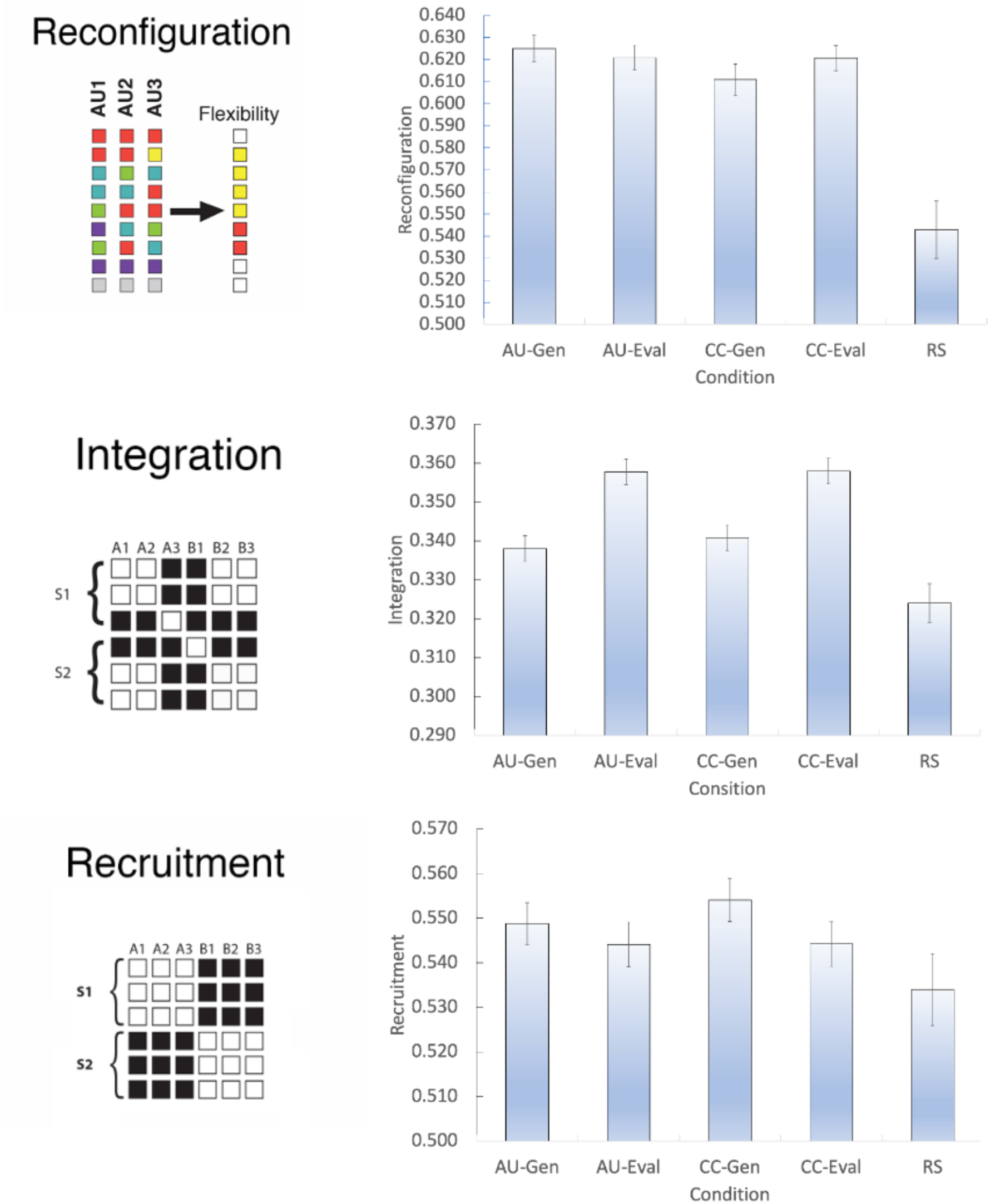
Whole-brain analysis of reconfiguration, integration, and recruitment across the four task-based conditions (Generation/Evaluation × AU/CC). In addition, these condition specific neural measures are compared to a baseline computed from participants’ resting-state fMRI data.

*Reconfiguration*: A Response Type (AU, CC) × Task (Generation, Evaluation) mixed model ANOVA was used to examine the effects of condition and time on whole-brain reconfiguration. This analysis revealed a significant main effect of Response Type, *F*(1, 41) = 4.29, *p* < .045, *η*^2^ = .095. Post-hoc independent-samples *t*-test analyses showed that this effect was driven by AU responses being associated with higher reconfiguration (*M* = .623, *SD* = .03) than CC responses (*M* = .616, *SD* = .03), *t*(41) = 2.07, *p* = .045, *d* = .32.

In addition, this analysis revealed a marginally significant interaction between Response Type and Task, *F*(1, 41) = 4.024, *p* = .051, *η*^2^ = .089. Post-hoc paired-samples *t*-test analyses showed that this effect was driven by a significant difference in the reconfiguration measure across the two conditions. A significantly higher reconfiguration measure was obtained for generating AU (*M* = .625, *SD* = .04) compared to CC (*M* = .611, *SD* = .05) responses, *t*(41) = 2.04, *p* = .047, *d* = .045. No significant differences were found in the reconfiguration measure between evaluating AU (*M* = .621, *SD* = .037) and CC (*M* = .621, *SD* = .037) responses, *t*(41) = 1.19, *p* = .24 *d* = .18.

*Integration*: A similar mixed ANOVA design was used to examine the effects of condition and time on whole-brain integration. This analysis revealed a significant main effect of Task, *F*(1, 41) = 20.16, *p* < .001, *η*^2^ = .33. Post-hoc paired-samples *t*-test analyses revealed that this effect was driven by a significant difference in whole-brain integration during the evaluation task compared to the generation task. This effect was found in both the AU (Generation: *M* = .34, *SD* = .02, Evaluation: *M* = .35, *SD* = .02, *t*[41] = 4.53, *p* < .001, *d* = .70), and the CC (Generation: *M* = .34, SD = .02, Evaluation: *M* = .36, *SD* = .02, *t*[41] = 3.95, *p* < .001, *d* = .61) conditions.

*Recruitment*: No significant main effects of Response Type, *F*(1, 41) = 1.61, *p* = .21, *η*^2^= .04, Type, *F*(1, 41) = 2.39, *p* = .13, *η*^2^ = .06, or interaction, *F*(1, 41) = 1.44, *p* = .24, *η*^2^ = .03, were observed.

Finally, we tested whether the functional connectivity patterns reported above were specific to the generation tasks we employed in this study, as opposed to a more general response generation effect independent of task requirements. Accordingly, we permuted, for each measure separately, the relation of condition label and neural scores for all participants. This process was reiterated 100 times, and a participant’s permuted condition score was computed by averaging across these 100 iterations. We then conducted similar statistical analyses as reported above on the permuted conditions scores. No effects remain significant based on these permutation processes (all *p*’s > .5), indicating that our significant results are specific to the task conditions.

### Comparison to resting-state baseline

To examine the extent of specific task condition (generating vs. evaluating of creative and non-creative ideas) on whole brain neural dynamics, we computed similar dynamic network measures (reconfiguration, integration, recruitment) on resting state (RS) fMRI data collected from the same participants (see *Methods* and Ref. ^36^). In this RS scan, participants were not presented with any external stimuli and conducted task-free mind wandering. We compared the whole-brain dynamic network measures of the whole-brain RS data with each of its corresponding measures for each of the task-based conditions using a paired-samples *t*-test (**Fig. 3**).

Whole-brain reconfiguration was lower for RS (mean = .543, SD = .085) than for the whole-brain reconfiguration of the four conditions: AU-Gen (mean = .625, SD = .039), *t*(41) = – 5.64, *p* < .001, *d* = .87; AU-Eval (mean = .621, SD = .037), *t*(41) = –6.20, *p* < .001, *d* = .96; CC-Gen (mean = .611, SD = .046), *t*(41) = –4.32, *p* < .001, *d* = .67; and CC-Eval (mean = .621, SD = .037), *t*(41) = –6.18, *p* < .001, *d* = .95.

Whole brain integration was lower for RS (mean = .324, SD = .032) than for the whole-brain integration of the four conditions: AU-Gen (mean = .338, SD = .022), *t*(41) = –2.58, *p* = .007, *d* = .40; AU-Eval (mean = .358, SD = .021), *t*(41) = –6.26, *p* < .001, *d* = .97; CC-Gen (mean = .341, SD = .021), *t*(41) = –3.07, *p* = .002, *d* = .47; and CC-Eval (mean = .358, SD = .021), *t*(41) = –6.30, *p* < .001, *d* = .97.

Finally, whole brain recruitment was significantly lower for RS (mean = .534, SD = .049) than for the whole-brain recruitment of the two generation conditions, and numerically, non-significantly lower compared to the two evaluation conditions: AU-Gen (mean = .549, SD = .031), *t*(41) = –1.93, *p* – .031, *d* = .30; AU-Eval (mean = .544, SD = .033), *t*(41) = –1.05, *p* = .15, *d* = .16; CC-Gen (mean = .544, SD = .032), *t*(41) = –2.20, *p* = .017, *d* = .35; and CC-Eval (mean = .544, SD = .032), *t*(41) = –1.06, *p* = .15, *d* = .16.

Overall, we find that participants’ RS fMRI is more stable (lower reconfiguration, integration, and recruitment), relative to the task-based conditions—indicating increased cross-system synchronization at rest (**Fig. 3**).

### Examining the task-specificity of the neural integration effect for evaluation

Next, we examined whether the neural integration effect for evaluation during creative thinking was task-specific, and not a broad-spectrum effect of generally evaluating whether one’s response is appropriate for any task. We did so by computing participants’ whole-brain neural measures of reconfiguration, integration, and recruitment during their evaluation of creative and non-creative responses generated by other participants in a previous study ^47^. Specifically, besides evaluating their own generated ideas, all participants evaluated the responses generated by external participants to the same 8 AU and 8 CC objects (see *Methods and Materials*).

These analyses did not reveal any significant differences in the neural measures of evaluation of other people’s ideas: Whole-brain reconfiguration for AU (mean = .36, SD = .04) was not significantly different than for CC (mean = .36, SD = .05), *t*(40) = –.64, *p* = .52, d = .1; whole-brain integration for AU (mean = .31, SD = .04) was not significantly different than for CC (mean = .31, SD = .04), *t*(40) = –.13, *p* = .90, d = .02; and whole-brain flexibility for AU (mean = .47, SD = .05) was not significantly different than for CC (mean = .46, SD = .05), *t*(40) = –.25, *p* = .81, d = .04.

### System level neural analysis

Finally, we conducted a similar analysis of neural reconfiguration, integration, and recruitment at the neural system level, focusing on the EN, DMN, and SN. To do so, we used the Yeo *et al.* method, which partitions the brain into 17 sub-systems ^48,49^. Based on our *a priori* predictions, we focused on three EN sub-networks: ConA (bilateral frontal and parietal regions), ConB (bilateral rostral and caudal frontal, inferior parietal and temporal regions), and ConC (bilateral precuneus); three DMN sub-networks: DefA (bilateral orbital superior frontal regions, IPL), DefB (bilateral superior frontal, mid-temporal regions), and DefC (bilateral hippocampus); and two SN sub-networks: SalA (bilateral superior frontal regions, insula), and SalB (bilateral rostral medial-frontal regions, left insula; see **Fig. 4**). For each of these subsystems, we averaged the three neural dynamic measures across all brain regions that comprise these systems. Finally, similar to the whole-brain analysis, we conducted a Response Type (AU, CC) × Task (Generation, Evaluation) mixed model ANOVA on the three measures of neural dynamics.

**Fig. 4.**
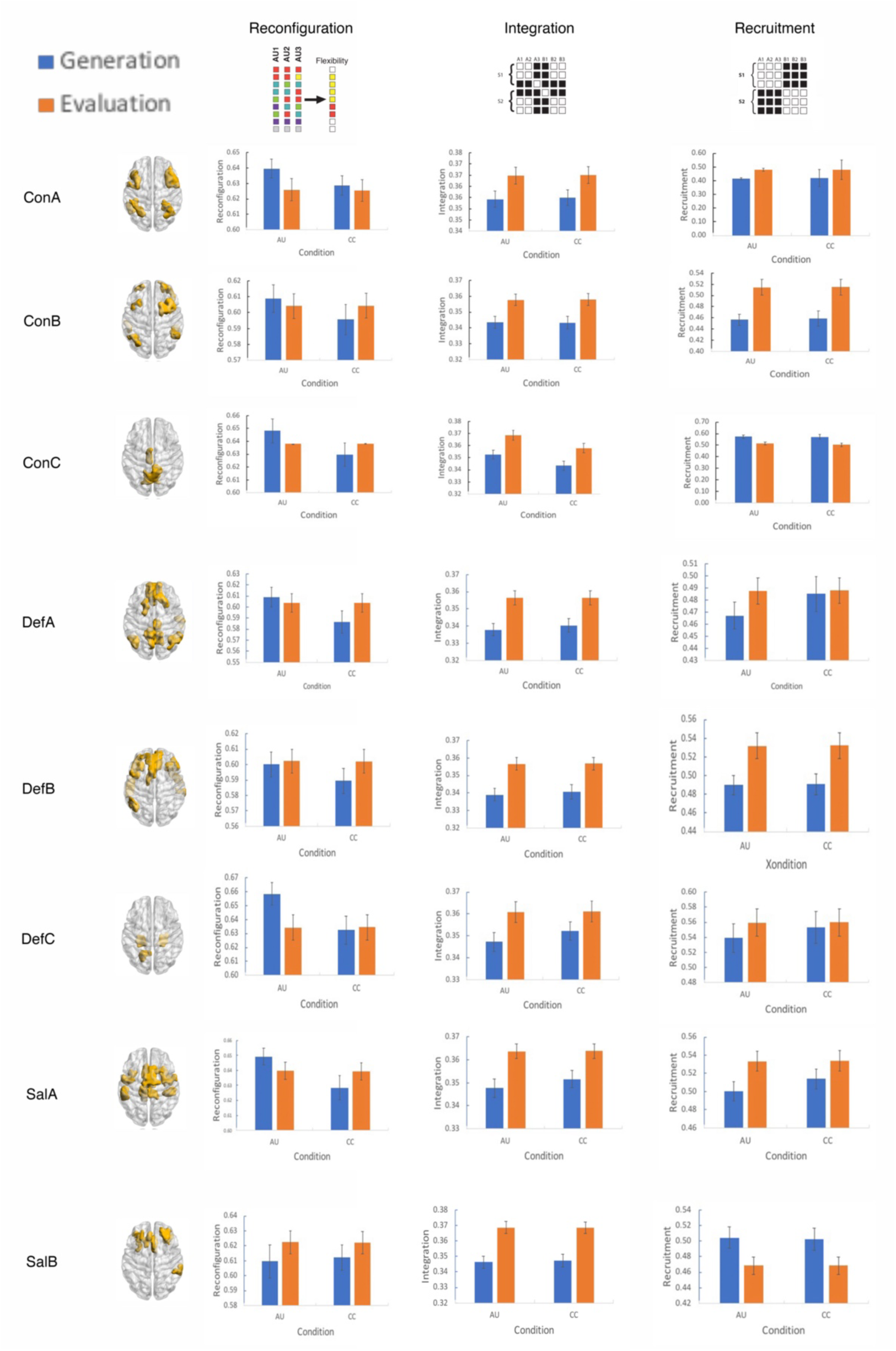
System-level brain analysis of neural reconfiguration (left), integration (center), and recruitment (right) across the four task-based conditions (Generation/Evaluation × AU/CC) for the seven neural systems analyzed. Neural systems are defined via the Yeo *et al.* partition of the brain into 17 sub-systems ^48,49^. We focused our analysis on three subnetworks of the EN: ConA (bilateral frontal and parietal regions), ConB (bilateral rostral and caudal frontal, inferior parietal and temporal regions), and ConC (bilateral precuneus); three subnetworks of the DMN: DefA (bilateral orbital superior frontal regions, IPL), DefB (bilateral superior frontal, mid-temporal regions), and DefC (bilateral hippocampus); and two subnetworks of the SN: SalA (bilateral superiorfrontal regions, insula), and SalB (bilateral rostral medial-frontal regions, left insula). Each neural system is illustrated via BrainNet Viewer ^50^.

*Reconfiguration*: A main effect of Response Type was found for the following systems (**Fig. 4**): DefA, *F*(1, 41) = 4.278, *p* = .045, *η*^2^ = .094, DefC, *F*(1, 41) = 7.596, *p* = .009, *η*^2^ = .156, and SalA, *F*(1, 41) = 5.712, *p* = .021, *η*^2^ = .123. In all these subsystems, reconfiguration was significantly higher for AU than for CC: DefA: *t*(41) = 2.068, *p* = .04, *d* = .32; DefC: *t*(41) = 2.756, *p* = .009, *d* = .43, and SalA: *t*(41) = 2.394, *p* = .021, *d* = .37.

In addition, a significant interaction effect between Response Type and Task was found for the same systems (**Fig. 4)**: DefA, *F*(1, 41) = 4.082, *p* = .05, *η*^2^ = .091, DefC, *F*(1, 41) = 8.179 *p* = .007, *η*^2^ = .16, and SalA, *F*(1, 41) = 5.404, *p* = .025, *η*^2^ = .110. Post-hoc *t*-tests revealed that this interaction effect was related to higher reconfiguration in generating AU compared to generating CC: DefA: *t*(41) = 2.045, *p* = .047, *d* = .32; DefC: *t*(41) = 2.81, *p* = .01, *d* = .43; and SalA: *t*(41) = 2.36, *p* = .023, *d* = .36.

*Integration*: A main effect of Task was found in the following systems (**Fig. 4)**: ConA, *F*(1, 41) = 5.76, *p* = .021, *η*^2^ = .123, ConB, *F*(1, 41) = 8.03, *p* = .007, *η*^2^ = .164, ConC, *F*(1, 41) = 10.761, *p* = .002, *η*^2^ = .208, DefA, *F*(1, 41) = 10.285, *p* = .002, *η*^2^ = .208, DefB, *F*(1, 41) = 13.743, *p* < .001, *η*^2^ = .251, DefC, *F*(1, 41) = 1.968, *p* = .053, *η*^2^ = .088, SalA, *F*(1, 41) = 13.748, *p* < .001, *η*^2^ = .251, and SalB, *F*(1, 41) = 21.148, *p* < .001, *η*^2^ = .340. Across all these systems, the Evaluation task was associated with significantly more integration compared to the Generation task: ConA: *t*(41) = –2.4, *p* = .021, *d* = .37; ConB: *t*(41) = –2.83, *p* = .007, *d* = .44; ConC: *t*(41) = – 3.28, *p* = .002, *d* = .51; DefA: *t*(41) = –3.28, *p* = .002, *d* = .51; DefB: *t*(41) = –3.71, *p* < .001, *d* = .57; DefC: *t*(41) = –1.99, *p* = .05, *d* = .31; SalA: *t*(41) = –3.71, *p* < .001, *d* = .57; and SalB: *t*(41) = –4.60, *p* < .001, *d* = .71. In addition, a main effect of Response Type was found for ConC, *F*(1, 41) = 12.943, *p* < .001, *η*^2^ = .240. Post-hoc *t*-test analysis revealed that this effect was due to higher integration for AU than CC, *t*[41] = 3.60, *p* < .001, *d* = .56.

*Recruitment*: A main effect of Task was found in the following systems (**Fig. 4**): ConA, *F*(1, 41) = 26.67, *p* = .001, *η*^2^ = .394, ConB, *F*(1, 41) = 16.223, *p* = .001, *η*^2^ = .284, ConC, *F*(1, 41) = 13.695, *p* = .001, *η*^2^ = .250, DefB, *F*(1, 41) = 8.118, *p* = .007, *η*^2^ = .165, and SalB, *F*(1, 41) = 5.046, *p* = .03, *η*^2^ = .110. For the ConA, ConB, and DefB systems, this effect was related to higher recruitment for Evaluation: ConA: *t*(41) = –5.17, *p* < .001, *d* = .80, ConB, *t*(41) = –4.03, *p* < .001, *d* = .62, and DefB, *t*(41) = –2.85, *p* = .01, *d* = .44. For the ConC and SalB systems, this effect was related to higher recruitment for Generation: ConC, *t*(41) = 3.70, *p* < .001, *d* = .57, and SalB, *t*(41) = 2.25, *p* = .03, *d* = .35.

## Discussion

Much recent work on the neuroscience of creativity has identified contributions of large-scale brain networks and their interactions to creative idea generation ^6,28,51^. A smaller number of studies has revealed similar contributions during one’s evaluation of the creativity of others’ ideas ^32,33^. However, the precise neural dynamics that support the generation and evaluation of creative ideas within the same person remain poorly understood (cf. ^32,52^). Although past research ^6,7,21,51^ has employed functional connectivity measures, these studies have generally relied on the average co-activation of regions across the brain, which is sub-optimal for precisely capturing critical, time-sensitive information on the dynamic contributions of particular brain regions during a creative task. Our study addresses this knowledge gap by means of a novel, within-subjects paradigm that allowed us to use network neuroscience methods to examine how large-scale networks interact during creative cognition.

Overall, our results showed that reconfiguration within default mode and salience network regions characterizes idea generation, but not evaluation, whereas large-scale system integration is a signature feature of idea evaluation, but not generation. These results are compatible with the prediction that the brain enters a state of transient instability during creative idea generation, possibly in support of the pursuit of novelty of the responses—a result further aligned with the behavioral differences in novelty between the two tasks (AU vs. CC). In contrast, the process of integration may reflect higher, general, collaboration across brain systems in the service of assessing both the novelty and the appropriateness of a response in context.

Our findings are an important contribution to the literature on the neural bases of creative cognition: our network neuroscience measures were able to capture not simply the co-engagement of different regions across the brain, but—importantly—the changes in connectivity both within and between systems during the different portions of the creative task and across time. Past work ^7,21^ has shown that, during creative idea generation, increased connectivity between EN and DMN, on average, is associated with performance. However, our approach demonstrates that this relationship is significantly more complex. Our tools show that what is critical is how the pattern of connections across these large-scale systems changes across time in support of creative behavior (see also Ref. ^52^). Specifically, our results reveal that for creative generation the interaction between response type and task for the reconfiguration measure was significant for the default mode and salience networks only, but not the executive network. This finding is inconsistent with past interpretations of the involvement of EN in creative cognition ^7^, and suggests that for creative generation the flexible engagement of systems related to memory retrieval is potentially more important than the engagement of systems involved in cognitive control (cf., Refs. ^9,15,44,47,53,54^).

Among the potential limitations of this work is the choice of the control, CC task, which elicited higher variability in responses and was more complex than other control tasks used in creativity neuroimaging research (e.g., Ref. ^21^). Participants generated either features or functions as common characteristics on this task, which led to potentially different assessment processes during the evaluation task. Yet, the integration measure revealed that the neural systems involved in evaluation processes are not exclusive to creative cognition, as they were also similarly engaged for the evaluation of responses in the CC task. Given this limitation, the specificity of the integration effect during the evaluation stage of creativity we report here needs to be interpreted with caution and would benefit from further study. However, previous work has shown how people overweigh novelty when evaluating creative responses over non-creative responses ^55,56^. Thus, although we cannot test this possibility directly, it is likely that participants in our study were focusing on novelty when evaluating the AU responses and on appropriateness when evaluating the CC trials. Finally, participants’ evaluation of ideas generated by other people—whether creative or non-creative—did not lead to the same integration effects found when they were evaluating their own ideas. This difference indicates that our neural integration effect for evaluation during creative thinking may be task-specific, and not a broad-spectrum effect of evaluating task performance more generally.

Another limitation is the block—and not event-related—design of our study. Collapsing condition specific trials together potentially adds a temporal confound to our findings and minimizes our ability to directly examine the neural dynamics of the creative process. Follow-up studies should replicate and extend our findings via an event-related design to better study the buildup of neural dynamics during the creative process, like previous studies examining neural reconfiguration in other cognitive tasks ^35,41^.

Overall, our results support the conclusion that creativity entails dynamic, parallel, and complex processes, involving multiple cognitive systems and their underlying neural mechanisms. Our study advances our understanding of this underlying complexity, by means of a unique within-subjects design and by applying dynamic network neuroscience methods. Such a design allows us to examine the neural bases of the prevalent generation-evaluation model of creative thinking ^57^, and the dynamics of how different functional neural systems interact to realize one of the most complex behaviors that humans evince ^6,7,21^.

## Methods

### Participants

Participants (*N* = 50) were recruited from the University of Pennsylvania. Five participants were excluded because they did not return for the second scan session. Two participants terminated the study due to nausea during the first scan session or due to becoming ineligible for MRI scanning after the first scan session. One participant was excluded due to poor performance on the task. As such, the final sample included 42 participants (26 females, mean age = 22.5 years [*SD* = 3.3], mean education = 16.4 years [*SD* = 2.51]). All participants were right-handed with normal or corrected-to-normal vision, and reported no history of neurological disorder, cognitive disability, or use of medication with potential to affect the central nervous system. Participants were monetarily compensated for their participation in the study. The study was approved by the University of Pennsylvania’s Institutional Review Board.

### Generation-evaluation task

The task consisted of two phases, a week apart, each conducted inside the scanner while participants underwent fMRI. In the Generation phase, participants were presented with 64 pictures of common objects ^58^. For half of these objects (*n* = 32), participants were asked to generate creative responses, namely, alternative uses for the objects (AU task); for the other half of the objects (*n* = 32), participants were asked to generate non-creative responses, namely common characteristics of the objects (CC task). A trial in the Generation phase began with a short fixation cross (500 ms) followed by a brief presentation of the object with an instruction above it to complete either the AU or CC task (2500 ms). For each trial, participants were subsequently required to generate verbally as many responses as they could for each object (15000 ms) before the next trial began (**Fig. 1A**). Participant’s responses during the generation task were audio-recorded, as well as manually typed concurrently by a research assistant.

A week later, participants came back to the scanner to complete the Evaluation phase. In the Evaluation phase, participants were presented with the same objects they saw during the Generation phase. Object task assignment (to the creative or non-creative task) also remained identical to the Generation phase. During the Evaluation phase, for each participant separately, each object was paired with three of the responses that specific participant gave for each of the objects: their first response, their final response, and an intermediate response. These three types of responses were chosen as to control for potential serial order confounds across participants ^59^. A trial in the Evaluation phase began with a short fixation cross (500 ms), followed by a short presentation of the object with the instruction above it, as presented in the Generation phase (2500 ms). Next, participants were presented with their three responses for that object (12000 ms). Participants were asked to evaluate these responses, and then verbally declare which of these three responses was the most novel and appropriate (3000 ms) before the next trial began (**Fig. 1A**). To control for any potential confounds arising from generating more verbal responses during the Generation phase compared to the Evaluation phase, for 50% of the trials, participants were required to ‘think aloud’ as they evaluated their responses following established procedures ^60,61^.

After participants evaluated all 64 objects to which they generated responses (AU and CC) in the generation phase, they underwent a final, general evaluation task. In this general evaluation task, participants were presented with an additional 16 objects, 8 with AU responses and 8 with CC responses. Participants saw the exact same responses for these 16 objects, taken from Ref. ^47^. This evaluation task allowed us to directly compare the neural dynamics related to evaluating ones’ own ideas compared to generally evaluating ideas that were generated by someone else. All presentation order and parameters of this final general evaluation task was identical to the main evaluation task as described above (including ‘thinking aloud’ for 50% of these additional objects).

### fMRI design

In accordance with Chai *et al.*, ^36^, trials were organized in a block design and were semi-randomly assigned into pairs of trials within a block (e.g., two AU trials followed by two CC trials). Participants completed 4 runs in both scan, each consisting of 4 experimental blocks and 4 fixation blocks, lasting 352 seconds. Experimental blocks lasted for 288 seconds (with 16 trials per block); fixation blocks lasting 16 seconds were interleaved between the experimental blocks. In both Generation and Evaluation phases, trials began immediately after the previous one ended. Condition order was counterbalanced across runs.

### SemDis scoring of participants’ creativity

To quantify participants creativity, we leveraged computational models of semantic distance. Semantic distance is a proxy of the novelty dimension of creative thinking—the extent to which an idea is conceptually distant from common ideas—by computing the similarity between concepts in large text-based corpora of natural language (e.g., textbooks). Recently, semantic distance in creativity research was validated by showing reliable and strongly positive correlations with subjective novelty ratings ^46^. This approach is based on an online website (*SemDis*; semdis.wlu.psu.edu) that computes semantic distance between cue words and participants’ open-ended responses. It utilizes five semantic models that have shown the highest correspondence with subjective originality ratings ^46^, thereby yielding semantic distance scores that are more generalizable for creativity measurement ^62^. For each participant, we computed the average semantic distance for all five models across all AU and CC trials, denoting how far (on average) their responses were in an averaged semantic model, from the original objects.

### MRI data acquisition and preprocessing

Magnetic resonance images were obtained using a 3.0 T Siemens Trio MRI scanner (Siemens Medical Systems, Erlangen, Germany) equipped with a 32-channel head coil. T1-weighted structural images of the whole brain were acquired on both Generation and Evaluation scans using a three-dimensional magnetization-prepared rapid acquisition gradient echo pulse sequence, repetition time (TR) = 1850 ms; echo time (TE) = 3.91 ms; voxel size = 0.9 mm × 0.9mm × 1 mm; flip angle = 8°; FoV = 240 mm. A field map was also acquired at each of the scan sessions, TR = 580 ms; TE 1 = 4.12 ms; TE2 = 6.52 ms; flip angle = 45°; voxel size = 3.0 mm × 3.0 mm × 3.0 mm; FoV = 240 mm, to correct geometric distortion caused by magnetic field inhomogeneity. In all resting-state and task-based scans, T2*-weighted images sensitive to blood oxygenation level-dependent contrasts were acquired using a slice accelerated multiband echo planar pulse sequence ^63,64^, TR = 500 ms; TE = 25 ms; flip angle = 45°; voxel size = 3.0 mm × 3.0 mm × 3.0 mm; FoV = 192 mm ^65^. The resting-state scan lasted 8 minutes with the exact same parameters. Both task-based scans were composed of 4 runs, each including 16 trials divided into 4 experimental blocks and 4 fixation blocks.

Preprocessing was performed via FSL ^66^ and FreeSurfer ^67^ through a suite of Matlab scripts, according to Ref. ^40^. Cortical reconstruction and volumetric segmentation of the anatomical data were performed with the FreeSurfer image analysis suite ^68^. Functional data were de-spiked by replacing voxel values greater than 7 RMSE from a 1-degree polynomial fit to the time course of each voxel with the average value of the adjacent TRs. Motion correction parameters were computed by registering each volume of each run to the middle volume using a robust registration algorithm (mri_robust_register; ^69^) and voxel shift maps for EPI distortion correction that were calculated using PRELUDE ^70^ and FUGUE ^71^. The resulting transformations were combined and simultaneously applied to the functional images. Boundary-based registration between structural and functional images was performed with *bbregister* ^72^. Nuisance time series signals were regressed from the preprocessed data. These nuisance regressors included: a) 24 motion regressors ^73^; b) the five first principal components of non-neural sources of noise, obtained with FreeSurfer segmentation tools and removed, following the anatomical CompCor method ^74^; and c) an estimate of a local source of noise, estimated by averaging signals derived from the white matter located within a 15 mm radius of each voxel, following the ANATICOR method ^75^. The data were then high-pass filtered with a cutoff frequency of 0.009 Hz ^76^.

### Functional connectivity network construction

Functional brain networks are constructed using a gray matter parcellation based on the Lausanne atlas ^77,78^. This brain atlas parcellates the brain into 234 regions covering the cortex and subcortical regions. In line with previous studies ^40,41,79,80^, functional connectivity between these brain regions was computed based on continuous wavelet coherence, which identified areas in time frequency space where two time series co-varied in the frequency band 0.06–0.12 Hz ^81^. This frequency band has previously been used to measure functional associations between low-frequency components of the fMRI signal and task-related functional connectivity ^41,42,65,79^. Wavelet coherence functional connectivity matrices were computed using the continuous cross wavelet transform developed by Grinsted, Moore, and Jevrejeva ^81^. We apply a continuous—and not discrete—wavelet transform (CWT) to provide additional sensitivity to time-varying dynamics across our four conditions. The CWT produces a connectivity value between each pair of brain regions for each TR, sampled across the frequency band 0.06–0.12 Hz ^81^. These connectivity values were averaged across the frequency range to generate an averaged time-varying connectivity value between each region pair. This procedure resulted in 234×234 weighted adjacency matrices for each TR, with coherence values bounded between 0 and 1 for each functional connection or network edge.

### Multilayer network construction

We constructed a dynamical functional brain network that represented neural dynamics during generation and evaluation of creative and non-creative ideas. For each participant and each run, for each task separately (Generation, Evaluation), we averaged all TR-based CWT adjacency matrices of a trial, to generate averaged AU– and CC-trial level CWT functional connectivity matrices. Next, we pooled together all AU– and CC-trial CWT functional connectivity matrices by concatenating all relevant trials one after the other, thus, only partially preserving the temporal sequence of the BOLD signal. Finally, we coupled these Response Type (AU, CC) × Task (Generation, Evaluation) specific functional connectivity matrices in a multilayer network ^35,43,79^. In these AU/CC × Generation/Evaluation multilayered networks, each layer represents a different trial, and each brain region is connected to itself in adjacent layers by an identity link. Although layer/trial durations are short (18 seconds for both Generation and Evaluation trial), conducting such a multilayer network analysis in short time windows has been shown to highlight individual differences ^39^.

### Dynamic community detection

We used dynamic community detection techniques ^43,82^ to extract groups of brain regions that are functionally connected with one another, and to characterize how they reconfigure, integrate, and are recruited over conditions ^35,79^. This was achieved by applying a data-driven community detection algorithm on the functional connectivity adjacency matrices ^83,84^. Intuitively, community detection techniques aim to categorize network nodes into communities or clusters. To do so, we maximize a quality function called the multilayer modularity Q, with the associated maximum of Q called the maximum modularity. The modularity quality function describes the partitioning of a network’s nodes into communities via a comparison to a statistical null model ^85^. High values of Q indicate that the nodes of the network can be partitioned sensibly into modules with similar BOLD activity. A generalization of the modularity quality function for multilayer networks can be written as:

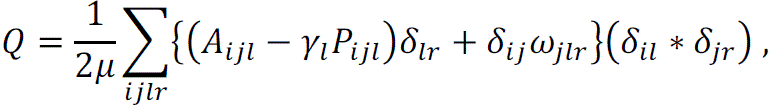

where *l* is the number of layers in the network, μ is the total edge weights in the network, *A_ijl_* is an adjacency matrix of a specific layer, P*_ijl_* is the corresponding null model of the layer ^86^, γ_l_ is a structural resolution parameter that defines the weights of intralayer connections, δ*_il_* denotes the community assignment of node *i* in layer *l*, δ*_jr_* denotes the community assignment of node *j* in layer *r*, and ω*_ilr_* denotes the connection strength between nodes across two layers (*l* and *r*). Importantly, changing the range of γ (number of communities) and ω (connection strength across layers) can affect the number and temporal dynamics of the detected communities ^40^. However, in order to not bias results toward a specific number of communities or a specific scale of temporal dynamics in community structure, we set γ and ω equal to one ^38,65^.

We optimized multilayer modularity using a generalization of a Louvain-like locally greedy algorithm ^87,88^ to yield a partition of regions into communities for each layer of each of the four multilayered networks. According to recent methodological recommendations ^82^, the Louvain algorithm was realized with the ‘moverandw’ as its randomization method. This randomization method leads to more reliable results of the Louvain community detection algorithm ^82^. Since the community detection algorithm is non-deterministic ^89^, we optimized the multilayer modularity quality function 100 times for each participant for each of the multilayered network ^35^. Finally, to resolve the variability across the 100 iterations of the community assignment partitions, we conduct a consensus analysis to identify the community assignment partition that summarizes the commonalities across the entire distribution of partitions for each one of the 32 layers separately ^90,91^. The results of this process are data-driven consensus-based identified communities for each of the 32 layers. Participants’ resting-state data was similarly analyzed by dividing the resting-state time series into 32 equal parts of 15 seconds.

### Neural dynamic measures

Across both task-based scans and resting-state scan, we computed for each participant and each condition three neural dynamic measures: Flexibility reconfiguration, integration, and recruitment. The *flexibility reconfiguration* of a node is defined as the probability that a node changes its community assignment across layers of the multilayer network ^79^. Since the slices in the multilayer networks convey different trials (AU or CC), we treat these networks as categorical, where any such community assignment change can occur across any pair of possible layers in the multilayer network. High values of flexibility indicate greater network reconfiguration.

The *integration* and *recruitment* of a node are calculated from a module allegiance matrix, which defines the percentage of layers in the multilayer network that node *i* and node *j* co-occur in the same community ^37^. To do so, each brain region was assigned to a resting-state based neural system, as defined by the Yeo *et al.* partition of the brain into 17 sub-systems ^48,49^ (see below).

The *integration* of a node is defined as the average probability that brain region *a* from brain system *b* will be assigned to the same community with other brain regions from other brain systems. High values of integration indicate greater cross-system interaction. The *recruitment* of a node is defined as the average probability that brain region *c* from brain system *d* is assigned to the same community with other brain regions from that brain system. High values of recruitment indicate greater brain system cohesiveness. Consistent with previous studies ^41,79,92^, we defined the reconfiguration, integration, and recruitment of the network over the entire brain as the mean score over all nodes in the network (*N* = 234). We then averaged these three neural dynamic measures across all participants, runs, and optimizations, to obtain representative measures for the entire group for each of the conditions (Generation/Evaluation × AU/CC).

### Resting state multilayer analysis

The resting-state multilayer network construction was conducted similarly to the task-based multilayer networks. The entire RS time-series was segmented into 32 equal time length parts, each of 18 seconds. Such RS time windows match the time length of each of the task-based layers in the task-based multilayer networks. To better equate the RS multilayer analysis to the task-based multilayer analysis (where trials were concatenated together), and to minimize the confound of time, we shuffled the order of the RS layers before conducting the multilayer analysis. A similar CWT approach was applied to arrive at a matched RS multilayer network for every participant. We reiterated the shuffling procedure of the RS layers and subsequent RS multilayer analysis 100 times. Whole-brain RS reconfiguration, integration, and recruitment measures were computed for each iteration. Finally, we computed the mean score for each measure over the 100 iterations.

### Brain network system level analysis

Given our predictions regarding the roles of DMN, EN, and SN in creativity ^6,21^, we also computed these three measures for specific brain networks. This goal is achieved via the Yeo *et al.* partition of the brain into 17 sub-systems ^48,49^. Thus, we focus our analysis, according to this partition, on the three subnetworks of the EN: ConA (bilateral frontal and parietal regions), ConB (bilateral rostral and caudal frontal, inferior parietal and temporal regions), and ConC (bilateral precuneus); three subnetworks of the DMN: DefA (bilateral orbital superior frontal regions, IPL), DefB (bilateral superior frontal, mid-temporal regions), and DefC (bilateral hippocampus); and two subnetworks of the SN: SalA (bilateral superiorfrontal regions, insula), and SalB (bilateral rostral medial-frontal regions, left insula).

### Procedure

The study consisted of two imaging sessions, a week apart. Prior to the first session, participants were screened for their ability to undergo fMRI scans, signed a consent form and were presented with the instructions for the Generation task that they were to complete during the first imaging session. During the first imaging session, each participant was presented again with the instructions of the Generation task and then placed in the MRI scanner. Padding around the head was used to minimize movement. Next, a high-resolution anatomical scan (2 minutes) and a resting-state (8 minutes) scan were collected. Following these scans, the participants completed the Generation task (23.46 minutes; as described above). The task began with a short practice, using objects that were not presented during the main task. Participants gave responses via an MRI compatible microphone (Optoacoustics, Inc. ®) and their responses were recorded and typed by the experimenter. After the scanning session, participants were taken out of the scanner and debriefed. During the second imaging session, the participant was presented with the instructions for the Evaluation task. Next the participant was placed in the MRI scanner with padding around the head to minimize movement. The scanning session began with a high-resolution anatomical scan followed by the Evaluation task (23.46 minutes; as described above). Finally, participants underwent a diffusion spectrography imaging scan (19 minutes, not analyzed in the current study). After the scanning session, participants were taken out of the scanner and debriefed.

## Acknowledgments

We thank Michelle Johnson and Mariya Bershad for their help in data collection. We also thank Richard Betzel, Arian Ashourvan, Lucy Chai, and Nathan Tardiff for fruitful discussions that led to the development of this study. This research was funded by an NIH award to S.T.-S. (R01 DC015359-02). D.S.B. also acknowledges support from the John D. and Catherine T. MacArthur Foundation, the Alfred P. Sloan Foundation, the ISI Foundation, the Paul Allen Foundation, the Army Research Laboratory (W911NF-10-2-0022), the Army Research Office (Bassett-W911NF-14-1-0679, Grafton-W911NF-16-1-0474, DCIST-W911NF-17-2-0181), the Office of Naval Research, the National Institute of Mental Health (2-R01-DC-009209-11, R01-MH112847, R01-MH107235, R21-M MH-106799), the National Institute of Child Health and Human Development (1R01HD086888-01), the National Institute of Neurological Disorders and Stroke (R01 NS099348), and the National Science Foundation (BCS-1441502, BCS-1430087, NSF PHY-1554488 and BCS-1631550). L.C. acknowledges support from the National Science Foundation Division of Research on Learning (NSF-DRL-2100137).

